# VIRTUS: a pipeline for comprehensive virus analysis from conventional RNA-seq data

**DOI:** 10.1101/2020.05.08.085308

**Authors:** Yoshiaki Yasumizu, Atsushi Hara, Shimon Sakaguchi, Naganari Ohkura

## Abstract

The possibility that RNA transcripts from clinical samples contain plenty of virus RNAs has not been pursued actively so far. We here developed a new tool for analyzing virus-transcribed mRNAs, not virus copy numbers, in the data of conventional and single-cell RNA-sequencing of human cells. Our pipeline, named VIRTUS (VIRal Transcript Usage Sensor), was able to detect 763 viruses including herpesviruses, retroviruses, and even SARS-CoV-2 (COVID-19), and quantify their transcripts in the sequence data. This tool thus enabled simultaneously detecting infected cells, the composition of multiple viruses within the cell, and the endogenous host gene expression profile of the cell. This bioinformatics method would be instrumental in addressing the possible effects of covertly infecting viruses on certain diseases and developing new treatments to target such viruses.

**Availability and implementation:** VIRTUS is implemented using Common Workflow Language and Docker under a CC-NC license. VIRTUS is freely available at https://github.com/yyoshiaki/VIRTUS.

**Supplementary information:** Supplementary data are available at Bioinformatics online.

## 1. Introduction

A variety of virus species including retroviruses, flaviviruses and herpesviruses, might contribute to the development of human diseases including autoimmune diseases and cancers. For example, Epstein-Barr virus (EBV) has been reported to play a causative role for head-neck cancer and lymphoma (Zapatka *et al*., 2020), and possibly for multiple sclerosis and systemic lupus erythematosus (Harley *et al*., 2018). It remains to be determined, however, which viruses are present in normal tissues and whether their state of activation contributes to disease development. Viruses can be detected by several methods such as antibody-based assays and PCR. Virus copy numbers in the genome can also be assessed by analyzing NGS derived data such as VirTect (Khan *et al*., 2019) and Kraken2 (Wood *et al*., 2019). On the other hand, it has been technologically difficult to examine the state of the virus in host tissues especially in relation to endogenous expression of host genes. In addition, since viral infection is heterogeneous depending on cell populations, it is unclear which cells are infected, how many virus species are present in the cells, and what states the viruses and the host cells assume. To address these issues, RNA information derived from polyA-based reverse transcription should be useful for analyzing intracellular viruses, since viruses intercept the host transcription systems, which yield polyA-tailed viral RNA transcripts along with endogenous RNAs from the host cells. We here attempted to establish a tool for measuring multiple viral transcriptomes even in a single cell.

## 2. VIRTUS

We developed a pipeline for detecting and quantifying transcripts of multiple viruses from conventional human RNA-seq data, and named it VIRTUS (VIRal Transcript Usage Sensor) (Supplementary Fig.1). As a framework of VIRTUS, RNA-seq data was quality-trimmed, filtered by fastp (Chen *et al*., 2018), and mapped to the human genome by STAR (Dobin *et al*., 2013). The unmapped reads were next aligned on 763 virus genome references. After removing polyX containing reads, infected viruses were determined comprehensively. Using salmon (Patro *et al*., 2017), a fixed amount of viral transcripts was quantified. The profiles of viral gene expression were integrated with the profiles of the host gene expression in each cell or sample.

## 3. Results

### 3.1. Application to bulk RNA-seq analyses

We first analyzed a bulk RNA-seq data of B cells infected with EBV (Mrozek-Gorska *et al*., 2019) (Fig.1 b,c). VIRTUS successfully detected EBV in all infected replicates (Supplementary Fig.3a); and the frequency of incorrect assignment of the virus infection was much less compared with other tools, such as VirTect and kraken2 (Supplementary Fig.3b-d). It was also able to quantify the EBV transcripts (Fig.1b, Supplementary Fig.3e) and detect its splicing pattern (Fig 1.c, Supplementary Fig.3h). We next evaluated virus contents in clinical samples (Rai *et al*., 2020) from peripheral blood leukocytes from 12 systemic lupus erythematosus patients and 4 healthy donors, and detected human papillomavirus 71,82; human herpesvirus 4, 5, 7; and human adenovirus C (Fig.1b). In addition, from bronchoalveolar lavage fluids from two SARS-CoV-2 infected patients, VIRTUS successfully detected SARS-CoV-2 in both patients (Supplementary Fig.4; Chen *et al*., 2020).

**Figure 1.**
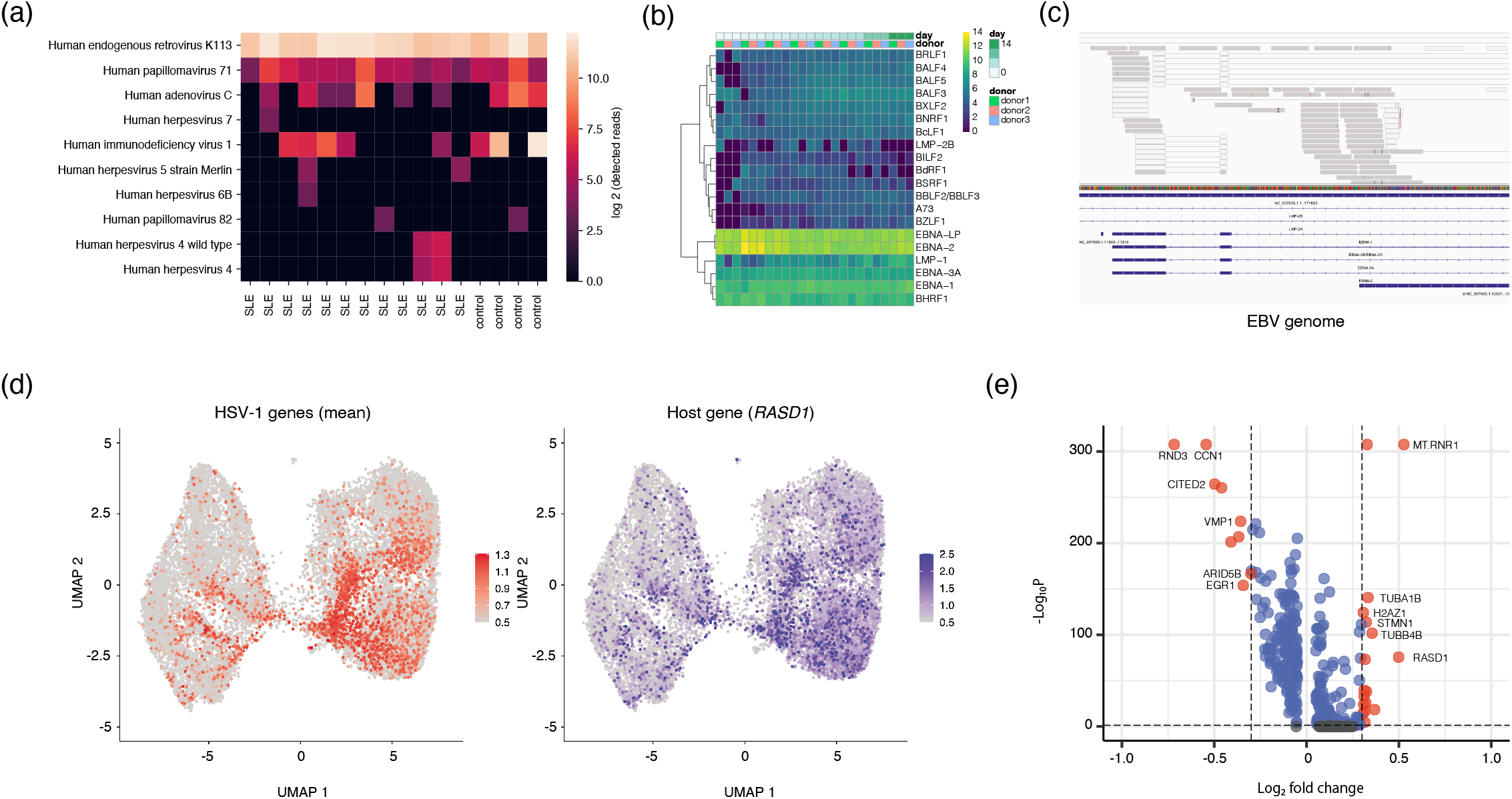
VIRTUS, a pipeline for analyzing multiple viruses, and its outputs from conventional RNA-seq data. (a) Viruses detected from peripheral blood leukocytes from SLE patients and healthy donors. (b) Top20 differentially expressed genes within EBV infected cells. (c) Virus-mapped reads visualized by The Integrative Genomics Viewer. (d) Mean transcripts of HSV-1 and the expression of a correlated host gene, *RASD1*, on UMAP plots. (e) Differentially expressed genes between HSV-1 infected and non-infected cells.

### 3.2. Application to single-cell RNA-seq analyses

We next applied VIRTUS to droplet-based single-cell RNA-seq data of human primary fibroblasts infected with Herpes simplex virus 1 (HSV-1) (Wyler *et al*., 2019) (Fig.1d,e). First, we conducted pooled screening of viruses, in which all reads from all cells were assigned at once, and detected HSV-1 in the samples. Then, we measured HSV-1 transcripts by Alevin (Srivastava *et al*., 2019), which was suitable for downstream analysis of VIRTUS. We detected infected single cells, and found differentially expressed genes, such as *RASD1* and *MT-RNR1*, between infected and non-infected cells, using VIRTUS and a standard single-cell pipeline. As shown in Fig. 1d and 1e, *RASD1*, one of the differentially expressed genes, was tightly linked to the HSV-1 infected cells.

## Conclusion

We developed a novel viral transcriptome detection and quantification pipeline, VIRTUS, which can be applied to both bulk and single-cell RNA-seq analyses. With this tool, we are able o detect the cells harboring activated viruses, the composition of multiple viruses in a cell, and the expression differences between infected and uninfected cells. It would help our understanding of how viruses contribute to certain diseases as a trigger or modifier of disease development and devising new ways of treatment by targeting viruses.

## Notes

https://github.com/yyoshiaki/VIRTUS

### Competing Interest Statement

The authors have declared no competing interest.

## References

Chen,L. et al. (2020) RNA based mNGS approach identifies a novel human coronavirus from two individual pneumonia cases in 2019 Wuhan outbreak. Emerging Microbes and Infections, 9, 313–319.

Chen,S. et al. (2018) Fastp: An ultra-fast all-in-one FASTQ preprocessor. In, Bioinformatics. Oxford University Press, pp. i884–i890.

Dobin,A. et al. (2013) STAR: Ultrafast universal RNA-seq aligner. Bioinformatics, 29, 15–21.

Harley,J.B. et al. (2018) Transcription factors operate across disease loci, with EBNA2 implicated in autoimmunity. Nature Genetics, 50, 699–707.

Khan,A. et al. (2019) Detection of human papillomavirus in cases of head and neck squamous cell carcinoma by RNA-seq and VirTect. Molecular Oncology, 13, 829–839.

Mrozek-Gorska,P. et al. (2019) Epstein–Barr virus reprograms human B lymphocytes immediately in the prelatent phase of infection. Proceedings of the National Academy of Sciences of the United States of America, 116, 16046–16055.

Patro,R. et al. (2017) Salmon provides fast and bias-aware quantification of transcript expression. Nature Methods, 14, 417–419.

Rai,R. et al. (2016) RNA-seq analysis reveals unique transcriptome signatures in systemic lupus erythematosus patients with distinct autoantibody specificities. PLoS One, 11, e0166312.

Srivastava,A. et al. (2019) Alevin efficiently estimates accurate gene abundances from dscRNA-seq data. Genome Biology, 20, 65.

Wood,D.E. et al. (2019) Improved metagenomic analysis with Kraken 2. Genome Biology, 20, 1–13.

Wyler,E. et al. (2019) Single-cell RNA-sequencing of herpes simplex virus 1-infected cells connects NRF2 activation to an antiviral program. Nature Communications, 10, 4878.

Zapatka,M. et al. (2020) The landscape of viral associations in human cancers. Nature Genetics, 52, 320–330.

