## Supplementary Data for "VIRTUS: a pipeline for comprehensive virus analysis from conventional RNA-seq data"

Contents

1. Overview of VIRTUS
   i. Pipeline description of VIRTUS
   ii. Comparison with other tools
2. Applications and methods
   i. Bulk RNA-seq from B cells infected with EBV
   ii. Bulk RNA-seq from peripheral blood leukocytes of SLE patients
   iii. Bulk RNA-seq from bronchoalveolar lavage fluids from two SARS-CoV-2 infected patients
   iv. Single-cell RNA-seq from primary fibroblasts infected with HSV-1
3. Code availability
4. References

#### 1. Description of the VIRTUS pipeline

We constructed a pipeline to detect and quantify virus transcripts comprehensively from conventional RNA-seq data of human specimens, and named it VIRTUS (Figure 1a, Supplementary Fig.1). In the pipeline, RNA-seq reads are trimmed and filtered by fastp with polyX filtering. Then, processed reads are aligned on the human genome by STAR. The unmapped reads are next aligned on the virus references consisted of 763 virus genome sequences (Supplementary Table 1). The references, which contain most viral genomes registered in RefSeq, are originated from VirTect, and updated based on the current virus diversity. For example, we added sequences of SARS-CoV-2 (NC_045512), a responsible virus for COVID-19, to the references. After filtration of polyX and low hit viruses (the default threshold is 400 reads), the summary of detected viruses is generated. Regarding the viruses whose transcripts were already determined, users can generate viral gene expression tables quantified by salmon with --validateMappings and --gcBias options. Users can install, create indices for VIRTUS, and run VIRTUS by one command.

#### ii. Comparison with other tools

VIRTUS, which we developed for compensating the insufficiencies of currently available NGS tools, has three unique characteristics. First, VIRTUS uses viral RNA transcripts as primary mapping targets so that the output reflects virus states, while most tools and analyses aim to detect viral genome copy numbers, and deal with the coverage of viral reads on the viral genome as the amounts of viruses. Second, measuring viral transcripts enabled us to assess viral states in a single-cell resolution. While no other virus detection tool had not been applied to single-cell RNA-seq data so far, we are able to detect multiple virus transcripts from a conventional single-cell RNA-seq data by VIRTUS. Third, VIRTUS adopted the recent technologies, such as Common Workflow Language (cwl; Amstutz *et al.*, 2016), Rabix Composer, and Docker, as its framework, for enabling easy installation and editing.

We compared VIRTUS to other virus detection tools. For virus detection, there are two strategies: alignment-based methods and k-mer assignment-based methods. Most alignment-based strategies have adopted tools with some disadvantages; for example, Tophat, which needs a long runtime, and bwa, which aligns reads on virus genomes without considering splicing events. On the other hand, k-mer based methods are relatively fast, but false-positivity is not negligible. To evaluate the accuracy, we prepared artificial fastq files that contained both 1,000,000 human genome reads and 1,000 EBV genome reads (E-MTAB-7805) with 10 replicates. For manipulation of fastq files, we used sambamba, samtools, and seqtk. We independently implemented VIRTUS (alignment-based method with STAR), VirTect (alignment-based method with Tophat), and Kraken2 (k-mer assignment) on 10 separate replicates. VIRTUS successfully detected EBV transcripts without any other incorrectly assigned viruses, while VirTect and Kraken2 detected several viruses as false positives (Supplementary Fig2a-c).

#### 2. Applications and methods

##### i. Bulk RNA-seq of B cells infected with EBV

We first validated and evaluated VIRTUS upon a bulk RNA-seq data which was derived from human B-cells artificially infected with Epstein-Barr virus (EBV). The raw fastq data (E-MTAB-7805) was subjected to VIRTUS with a default parameter (VIRTUS.PE.cwl), and aggregated as EBV gene expression profiles using tximport. EBV transcript levels, which was represented as a proportion of EBV-mapped reads per all mapped reads, showed the highest one day after infection in all replicates (Supplementary Fig. 3a). In addition to EBV, we also detected a small amount of Human herpesvirus 7, Human papillomavirus 71, Hepatitis virus B, Simian virus 40, and human endogenous retrovirus K113, in the samples. We next compared VIRTUS to other virus analyzing tools, such as VirTect (with default parameters) and Kraken2 (with the viral library and default parameters). The two tools detected a large number of virus species compared to VIRTUS (Supplementary Fig. 3b,c). VirTect tended to confusingly detect RNA viruses with a polyA tail. Kraken2 showed many false positive viruses due to its library, which contains multiple species of viruses including those not infecting humans. In addition, VIRTUS succeeded in quantifying the transcripts of EBV and revealed complicated EBV gene regulation patterns after infection (Supplementary Fig. 3e). To evaluate the sensitivity of detecting viral transcripts, we measured the percentage of reads mapped on the exon on the EBV genome. Although EBV exon regions occupy 16% of the genome, the percentage of reads mapped on the exons was above 60% in all the samples (Supplementary Fig. 3f). After calculating the overlap of the reads with bedtools coverage, we selected the top 20 highest expressed genes among all samples and analyzed their virus gene coverage patterns using geneBody_coverage.py included in RSeQC (Supplementary Fig. 3g). We also observed viral splicing patterns from the VIRTUS outputs on the Integrative Genomics Viewer (IGV, Supplementary Fig. 3h).

ii. Bulk RNA-seq from peripheral blood leukocytes of SLE patients

To measure the performance of VIRTUS on clinical samples, we performed the VIRTUS (VIRTUS.PE.cwl) on RNA-seq data from peripheral blood leukocytes from 12 systemic lupus erythematosus (SLE) patients and 4 healthy donors (PRJNA318253). Detailed information of the samples is available in the original article (Rai *et al.*, 2016). We processed the raw fastq files by VIRTUS.PE.cwl with an option of the mapping cutoff (--hit_cutoff 10) to increase detection sensitivity, and plotted the detected viruses using Seaborn (Fig. 1a). In SLE patients, we detected human endogenous retrovirus K113, human adenovirus C, human herpesvirus 4, 5, 6B, 7, and HPV 71. While there were no statistical significances in differentially expressed viruses between SLE patients and healthy donors (Mann-Whitney U-test), we could detect multiple virus transcripts quantitatively in both. The significance would be improved by increasing the sample size. Thus it would be useful for conducting a comprehensive study on virus association with human diseases.

iii. Bulk RNA-seq from bronchoalveolar lavage fluids from two SARS-CoV-2 infected patients

To detect SARS-CoV-2 from clinical samples, we applied VIRTUS.SE.cwl on bronchoalveolar lavage fluid (BALF) from two SARS-CoV-2 positive patients (PRJNA601736). Note that RNA was not polyA-selected and sequence reads mapped on the human genome were removed by the authors. The raw fastq files were processed by

VIRTUS with the default parameters (VIRTUS.SE.cwl). In both samples, VIRTUS detected SARS-CoV-2 (NC_045512.2) only (Supplementary Fig.4a). To validate the specificity, we re-aligned the unmapped reads on the SARS-CoV-2 by HISAT2 with the default parameter and visualized them by IGV (Supplementary Fig.4b). We observed properly aligned reads over the whole genome regions. These results suggest that VIRTUS can be applicable to SARS-CoV-2 investigation.

iv. Single-cell RNA-seq from primary fibroblasts infected with HSV-1

It is expected that the mode of viral infection and the composition of viruses are highly diverse in each cell. To interpret the diversity of viral infection under single-cell resolution, we developed an analytical flow of VIRTUS for single-cell RNA-seq data. There are two strategies for VIRTUS to deal with single-cell RNA-seq data. First, for technologies in which fastq files are generated for each individual cell such as SmartSeq2, users can simply apply VIRTUS on each cell. Second, in the case of the method which creates a single fastq file for one experiment with UMI identifiers such as 10x Chromium and DropSeq, we recommend users to utilize VIRTUS.SE.cwl on a pooled fastq file, which is an original file before demultiplexing cells. After screening viruses contained in the whole single-cell RNA-seq data to create indices of the viruses (createindex_singlevirus.cwl), users can determine the transcript levels of the viruses upon each cell using Alevin, which is a fast and accurate quantification tool designed for single-cell RNA-seq.

As an example of this workflow, we examined DropSeq derived single-cell RNA-seq data from HSV-1 infected human primary fibroblasts (GSE123782). We evaluated the number of viral transcripts in each batch by utilizing VIRTUS.SE.cwl on the second read files of the paired-end fastq file. We first generated virus indices by VIRTUS (createindex_singlevirus.cwl) with the HSV-1 reference transcripts (NC_001806.2), and applied them to Alevin to quantify the transcripts of host cells and HSV-1. The sum of HSV-1 gene expression quantified by Alevin and HSV-1 reads detected by VIRTUS were correlated (Supplementary Fig.5a). We detected Herpes simplex 1 (HSV-1), human herpesvirus 2 and human endogenous retrovirus K113 in the single-cell RNA-seq data

(Supplementary Fig. 5b). To determine infected cells, we calculated the distribution of averaged HSV-1 gene expression for each individual cell (Supplementary Fig. 5c). Then, we determined the threshold for the infected cells to 0.5 based on the distribution (Supplementary Fig. 5d). We also processed the host gene expression of each cell by Seurat functions (NormalizeData, FindVariableFeatures, and ScaleData), and removed the batch effect by IntegrateData function. Lastly, we conducted dimensionality reduction of the data by UMAP (Supplementary Fig. 5d), calculated statistics of expressions by FindMarkers, and visualized them by an R package EnhancedVolcano (Fig.1e). We detected several genes synchronously expressed with HSV-1 infection, such as *RASD1* (Fig.1e) *TUBA1B*, and *H2AZ* (Supplementary Fig. 5e). In addition, the two most associated genes with the HSV-1 infection were *MT-RNR1* and *MT-ATP6* (Supplementary Fig. 5e), which are coded from mitochondrial DNA. This observation suggests that the expression of mitochondrial genes is affected by a viral infection, and provides a clue to address the pathogenesis of mitochondrial gene-associated diseases. For analyzing biological effects of viruses, VIRTUS would be useful since it can detect virus-infected cells and the host responses simultaneously.

#### 1. Code availability

VIRTUS is available at https://github.com/yyoshiaki/VIRTUS under a CC-NC license. All codes are available at https://github.com/yyoshiaki/VIRTUS_Manuscript .

#### 2. References

Peter Amstutz, Michael R. Crusoe, Nebojša Tijanić (editors), Brad Chapman, John Chilton, Michael Heuer, Andrey Kartashov, Dan Leehr, Hervé Ménager, Maya Nedeljkovich, Matt Scales, Stian Soiland-Reyes,L.S. (2016) Common Workflow Language, v1.0.

Rai,R. *et al.* (2016) RNA-seq analysis reveals unique transcriptome signatures in systemic lupus erythematosus patients with distinct autoantibody specificities. *PLoS One*, **11**, e0166312.

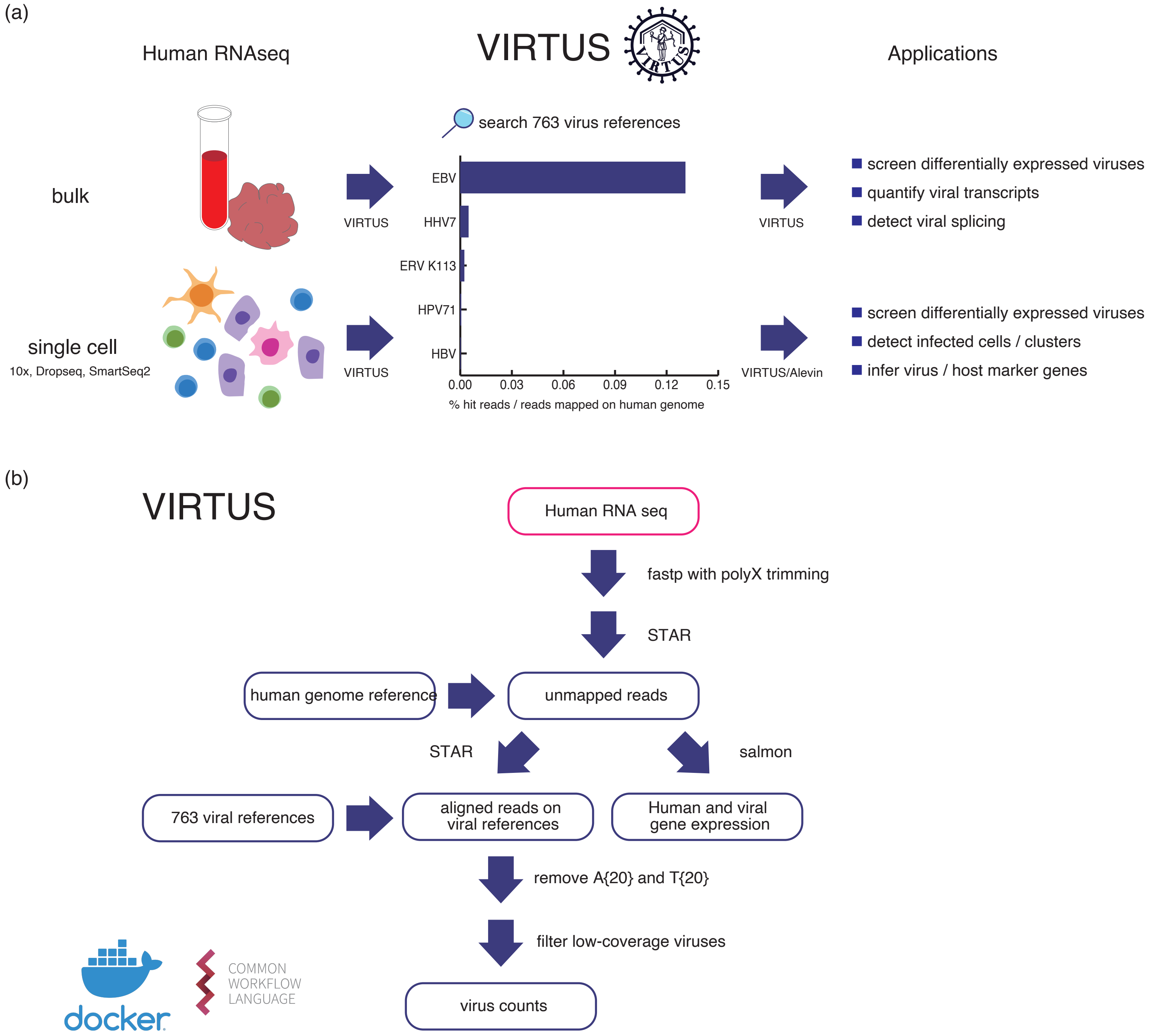


Supplementary Figure 1

Supplementary figure 1. VIRTUS schematic views of analytical flows (a) and the pipeline structure (b).

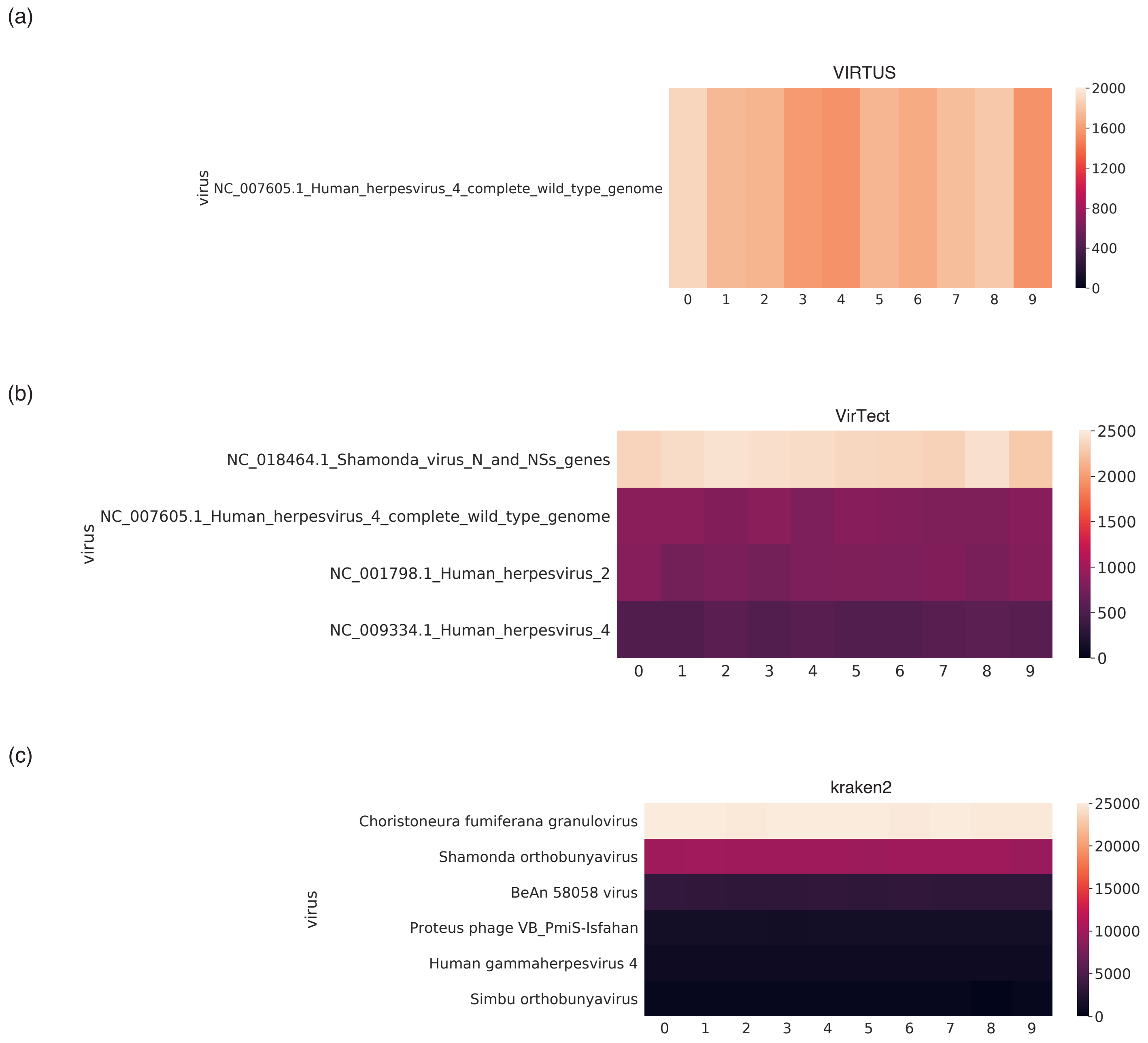


Supplementary Figure 2

Supplementary figure 2. Tool comparison by human reads with artificially mixed viral reads. 1,000 reads from EBV were mixed in 1,000,000 human mapped reads. The heatmaps are showing the number of detected viral transcripts or copies by VIRTUS (a), VirTect (b), and Kraken2 (c).

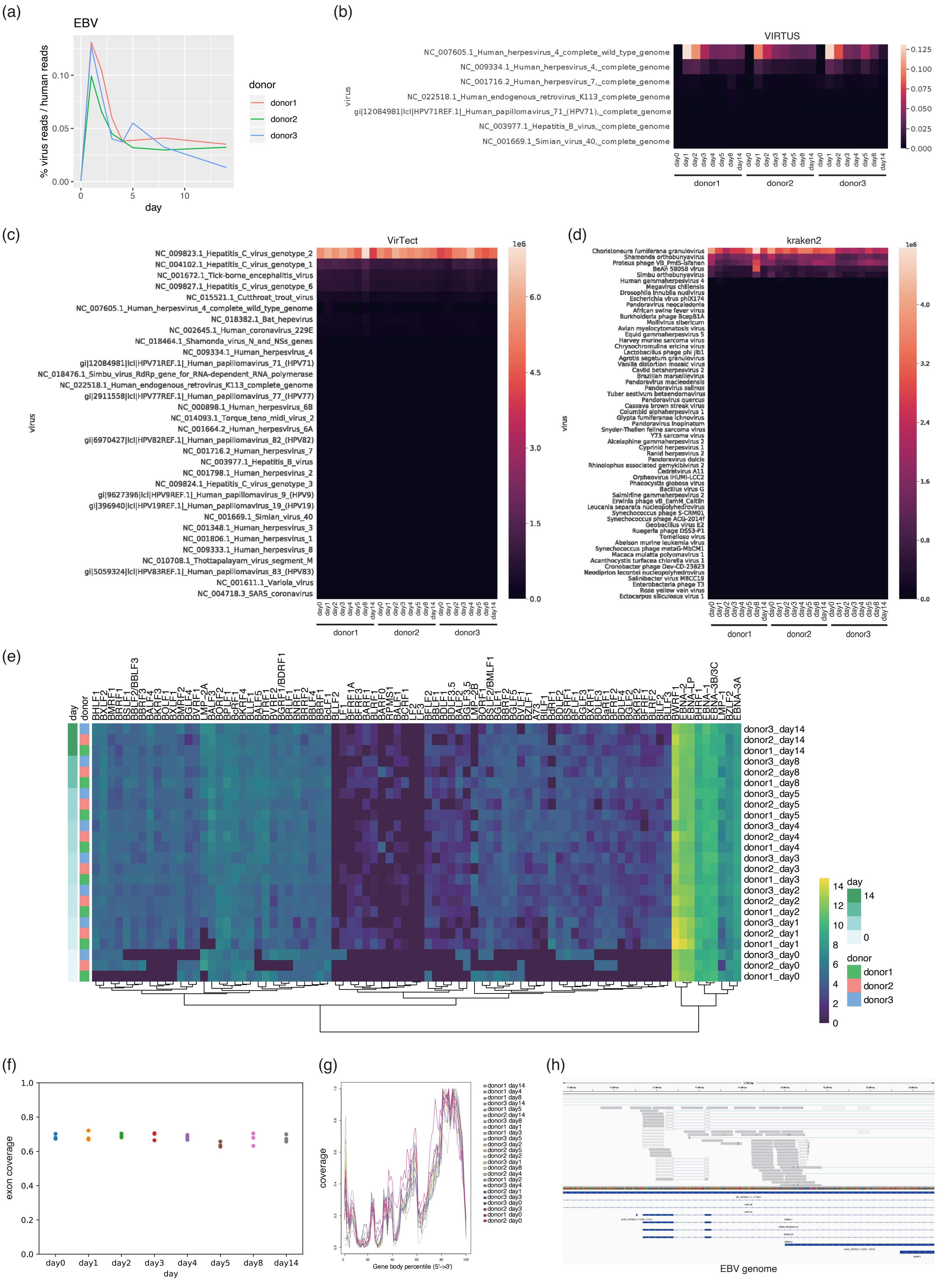


### Supplementary Figure 3

Supplementary figure 3. VIRTUS was validated by an EBV infection model. (a) The proportion of the reads which VIRTUS detected as EBV derived. The number of reads were normalized by the number of reads mapped on the human genome. (b-d) The proportion of the reads of all the viruses which are detected by three tools; VIRTUS (b), VirTect (c), and kraken2(d). The detected viral reads were normalized by the number of reads mapped on the human genome. (e) All EBV gene expression. (f) The percentage of reads mapped on the exon. (g) Gene body coverages for top20 highest expressed genes on EBV genome. (h) Mapped transcripts on the EBV genome.

(a)

| SRA RUN ID (patients) | Mapped reads on SARS-CoV-2 |
|---|---|
| SRR10903401 | 27342 |
| SRR10903402 | 107388 |

(b)

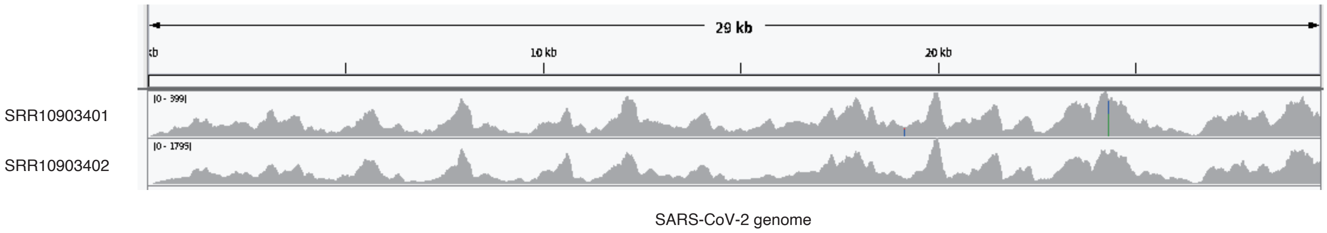


Supplementary Figure 4

Supplementary figure 4. The results of COVID-19 patients. (a) The number of mapped reads on the SARS-CoV-2 genome from BALF of two patients. (b)Aligned reads on SARS-Cov-2 visualized by IGV.

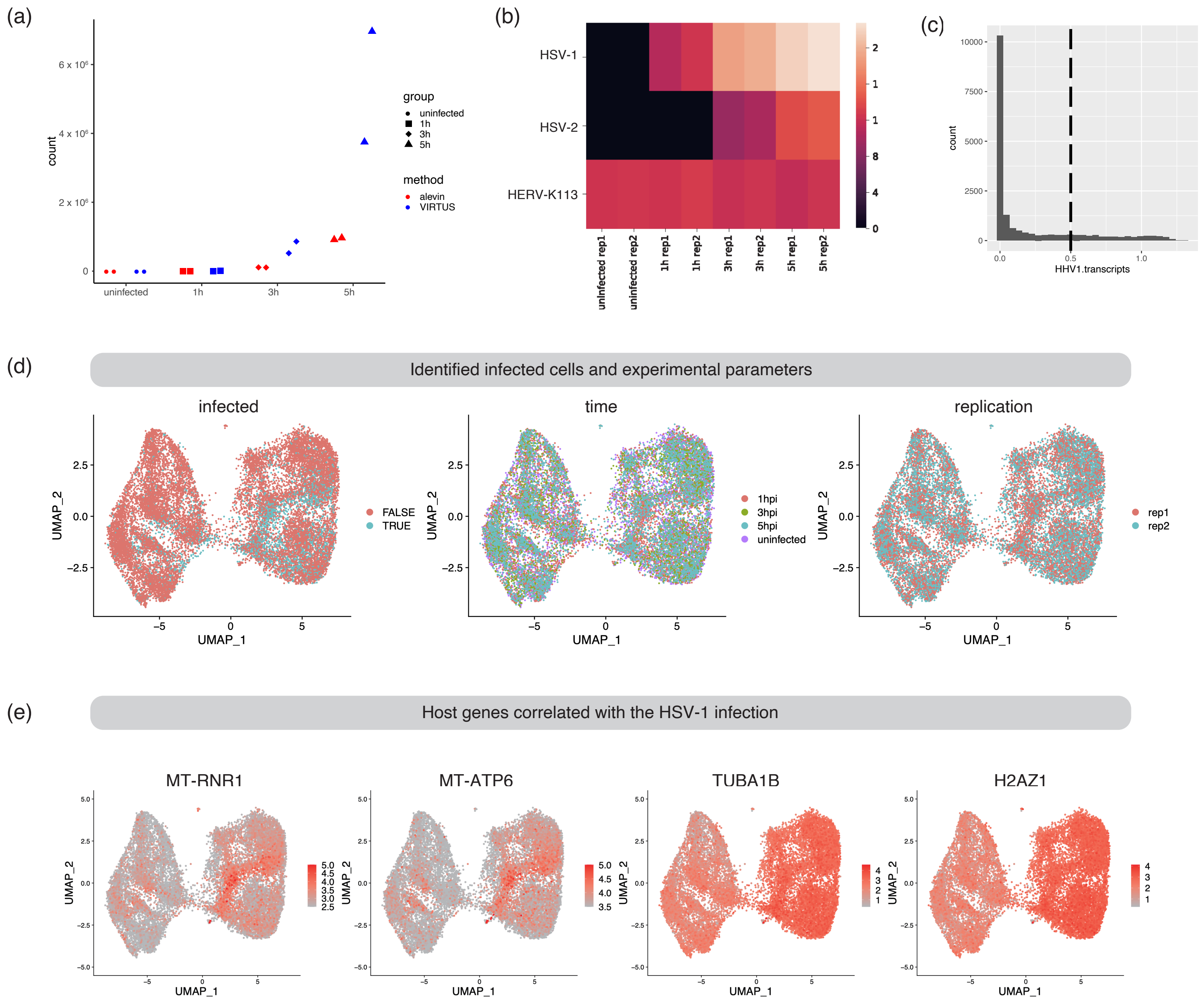


### Supplementary Figure 5

Supplementary figure 5. VIRTUS application on single-cell RNA-seq data from HSV-1 infected human primary fibroblasts. (a) The sum of HSV-1 gene expression quantified by Alevin and HSV-1 reads detected by VIRTUS. (b) All detected viruses by VIRTUS are shown. The color is $\log_2$ (the number of detected reads) (c) The distribution of mean gene expression of HSV-1. The dashed line is a threshold of infected cells. (d) Infected cells (left), time after the infection (center), and replicates (right) were shown on the UMAP plot. Each dot is every single cell.

### Supplementary Table 1. List of  enrolled viruses

| | | | |
|---|---|---|---|
| Abelson murine leukemia virus | Colorado tick fever virus segment 7 | Human enterovirus D | Human papillomavirus 27 |
| Adeno-associated virus - 1 | Colorado tick fever virus segment 8 | Human erythrovirus V9 | Human papillomavirus 28 |
| Adeno-associated virus - 2 | Colorado tick fever virus segment 9 | Human herpesvirus 1 | Human papillomavirus 29 |
| Adeno-associated virus - 3 | Cote d'Ivoire ebolavirus | Human herpesvirus 2 | Human papillomavirus 3 |
| Adeno-associated virus - 4 | Crimean-Congo hemorrhagic fever virus segment L | Human herpesvirus 3 | Human papillomavirus 30 |
| Adeno-associated virus - 7 | Crimean-Congo hemorrhagic fever virus segment M | Human herpesvirus 4 Type 1 | Human papillomavirus 31 |
| Adeno-associated virus - 8 | Crimean-Congo hemorrhagic fever virus segment S | Human herpesvirus 4 Type 2 | Human papillomavirus 32 |
| Adeno-associated virus 5 | Culex flavivirus | Human herpesvirus 5 strain Merlin | Human papillomavirus 33 |
| Adeno-associated virus 6 | Cupixi virus segment L | Human herpesvirus 6A | Human papillomavirus 34 |
| Adult diarrheal rotavirus strain J19 | Cupixi virus segment S | Human herpesvirus 6B | Human papillomavirus 35 |
| Adult diarrheal rotavirus strain J19 | Cutthroat trout virus | Human herpesvirus 7 | Human papillomavirus 36 |
| Adult diarrheal rotavirus strain J19 | Dengue virus 1 | Human herpesvirus 8 | Human papillomavirus 37 |
| Adult diarrheal rotavirus strain J19 | Dengue virus 2 | Human immunodeficiency virus 1 | Human papillomavirus 38 |
| Adult diarrheal rotavirus strain J19 | Dengue virus 3 | Human immunodeficiency virus 2 | Human papillomavirus 39 |
| Adult diarrheal rotavirus strain J19 | Dengue virus 4 | Human metapneumovirus | Human papillomavirus 4 |
| Adult diarrheal rotavirus strain J19 | Dobrava virus segment M | Human papillomavirus 1 | Human papillomavirus 40 |
| Adult diarrheal rotavirus strain J19 | Dobrava virus segment S | Human papillomavirus 10 | Human papillomavirus 41 |
| Adult diarrheal rotavirus strain J19 | Dobrava-Belgrade virus strain DOBV/Ano-Poroia/Afl9/1999 | Human papillomavirus 100 | Human papillomavirus 42 |
| Adult diarrheal rotavirus strain J19 | Dolphin morbillivirus | Human papillomavirus 101 | Human papillomavirus 43 |
| Adult diarrheal rotavirus strain J19 | Donggang virus | Human papillomavirus 102 | Human papillomavirus 44 |
| Aedes flavivirus | Dugbe virus segment L | Human papillomavirus 103 | Human papillomavirus 45 |
| Aguacate virus segment L | Dugbe virus segment M | Human papillomavirus 104 | Human papillomavirus 47 |
| Aguacate virus segment M | Dugbe virus segment S | Human papillomavirus 105 | Human papillomavirus 48 |
| Aguacate virus segment S | Duvenhage virus isolate 86132SA | Human papillomavirus 106 | Human papillomavirus 49 |
| Aino virus Gn-Gc-NSm gene for M polyprotein segment M genomic RNA isolate 38K | Ebola virus - Mayinga Zaire 1976 strain Mayinga | Human papillomavirus 107 | Human papillomavirus 5 |
| Aino virus N and NSs genes segment S genomic RNA isolate 38K | Entebbe bat virus | Human papillomavirus 108 | Human papillomavirus 50 |
| Aino virus RdRp gene for RNA-dependent RNA polymerase segment L genomic RNA isolate 38K | European bat lyssavirus 1 | Human papillomavirus 109 | Human papillomavirus 51 |
| Akabane virus segment L | European bat lyssavirus 2 | Human papillomavirus 11 | Human papillomavirus 52 |
| Akabane virus segment M | Eyach virus segment 1 | Human papillomavirus 110 | Human papillomavirus 53 |
| Akabane virus segment S | Eyach virus segment 10 | Human papillomavirus 111 | Human papillomavirus 54 |
| Alkhurma virus | Eyach virus segment 11 | Human papillomavirus 112 | Human papillomavirus 56 |
| Allpahuayo virus segment L | Eyach virus segment 12 | Human papillomavirus 113 | Human papillomavirus 57 |
| Allpahuayo virus segment S | Eyach virus segment 2 | Human papillomavirus 114 | Human papillomavirus 58 |
| Amapari virus segment L | Eyach virus segment 3 | Human papillomavirus 115 | Human papillomavirus 59 |
| Amapari virus segment S | Eyach virus segment 4 | Human papillomavirus 116 | Human papillomavirus 6 |
| Andes virus segment L | Eyach virus segment 5 | Human papillomavirus 117 | Human papillomavirus 60 |
| Andes virus segment M | Eyach virus segment 6 | Human papillomavirus 118 | Human papillomavirus 61 |
| Andes virus segment S | Eyach virus segment 7 | Human papillomavirus 119 | Human papillomavirus 62 |
| Apoi virus genome | Eyach virus segment 8 | Human papillomavirus 12 | Human papillomavirus 63 |
| Aravan virus | Eyach virus segment 9 | Human papillomavirus 120 | Human papillomavirus 65 |
| Arumowot virus segment L | Fer-de-lance virus | Human papillomavirus 121 | Human papillomavirus 66 |
| Arumowot virus segment M | Flexal virus segment L | Human papillomavirus 122 | Human papillomavirus 67 |
| Arumowot virus segment S | Flexal virus segment S | Human papillomavirus 123 | Human papillomavirus 68 |
| Australian bat lyssavirus | Friend murine leukemia virus | Human papillomavirus 124 | Human papillomavirus 69 |
| Avian metapneumovirus | GB virus C/Hepatitis G virus | Human papillomavirus 125 | Human papillomavirus 7 |
| Avian paramyxovirus 4 strain APMV-4/duck/Delaware/549227/2010 | Golden Gate virus segment L | Human papillomavirus 126 | Human papillomavirus 70 |
| Avian paramyxovirus 6 | Golden Gate virus segment S | Human papillomavirus 127 | Human papillomavirus 71 |
| BK polyomavirus | Goose paramyxovirus SF02 | Human papillomavirus 128 | Human papillomavirus 72 |
| Bagaza virus | Great Island virus segment 1 | Human papillomavirus 129 | Human papillomavirus 73 |
| Banna virus segment 1 | Great Island virus segment 10 | Human papillomavirus 13 | Human papillomavirus 74 |
| Banna virus segment 10 | Great Island virus segment 2 | Human papillomavirus 130 | Human papillomavirus 75 |
| Banna virus segment 11 | Great Island virus segment 3 | Human papillomavirus 131 | Human papillomavirus 76 |
| Banna virus segment 12 | Great Island virus segment 4 | Human papillomavirus 132 | Human papillomavirus 77 |
| Banna virus segment 2 | Great Island virus segment 5 | Human papillomavirus 133 | Human papillomavirus 8 |
| Banna virus segment 3 | Great Island virus segment 6 | Human papillomavirus 134 | Human papillomavirus 80 |
| Banna virus segment 4 | Great Island virus segment 7 | Human papillomavirus 135 | Human papillomavirus 81 |
| Banna virus segment 5 | Great Island virus segment 8 | Human papillomavirus 136 | Human papillomavirus 82 |
| Banna virus segment 6 | Great Island virus segment 9 | Human papillomavirus 137 | Human papillomavirus 83 |
| Banna virus segment 7 | Guanarito virus segment L | Human papillomavirus 138 | Human papillomavirus 84 |
| Banna virus segment 8 | Guanarito virus segment S | Human papillomavirus 139 | Human papillomavirus 85 |
| Banna virus segment 9 | Hantaan virus | Human papillomavirus 14 | Human papillomavirus 86 |
| Bat hepevirus | Hantaan virus | Human papillomavirus 140 | Human papillomavirus 87 |
| Bat sapovirus TLC58 HK | Hantaan virus segment L | Human papillomavirus 141 | Human papillomavirus 88 |
| Bear Canyon virus segment L | Hantavirus Z10 chromosome L | Human papillomavirus 142 | Human papillomavirus 89 |
| Bear Canyon virus segment S | Hantavirus Z10 chromosome S segment | Human papillomavirus 143 | Human papillomavirus 9 |
| Beilong virus | Hantavirus Z10 segment M | Human papillomavirus 144 | Human papillomavirus 90 |
| Borna disease virus | Hendra virus | Human papillomavirus 145 | Human papillomavirus 91 |
| Bovine parainfluenza virus 3 | Hepatitis B virus | Human papillomavirus 146 | Human papillomavirus 92 |
| Bovine respiratory syncytial virus | Hepatitis C virus genotype 1 | Human papillomavirus 147 | Human papillomavirus 93 |
| Brazoran virus segment L | Hepatitis C virus genotype 2 | Human papillomavirus 148 | Human papillomavirus 94 |
| Brazoran virus segment M | Hepatitis C virus genotype 3 genome | Human papillomavirus 149 | Human papillomavirus 95 |
| Brazoran virus segment S | Hepatitis C virus genotype 4 genome | Human papillomavirus 15 | Human papillomavirus 96 |
| Bundibugyo ebolavirus | Hepatitis C virus genotype 5 genome | Human papillomavirus 150 | Human papillomavirus 97 |
| Bunyamwera virus L segment | Hepatitis C virus genotype 6 | Human papillomavirus 151 | Human papillomavirus 98 |
| Bunyamwera virus M segment | Hepatitis E virus | Human papillomavirus 153 | Human papillomavirus 99 |
| Bunyamwera virus segment S | Hepatitis delta virus | Human papillomavirus 154 | Human parainfluenza virus 1 |
| Bussuquara virus | Human T-lymphotropic virus 1 | Human papillomavirus 155 | Human parainfluenza virus 2 |
| CAS virus segment L | Human T-lymphotropic virus 2 | Human papillomavirus 156 | Human parainfluenza virus 3 |
| CAS virus segment S | Human T-lymphotropic virus 4 | Human papillomavirus 159 | Human parainfluenza virus 4a viral cRNA strain: M-25 |
| California sea lion anellovirus | Human adenovirus 54 | Human papillomavirus 16 | Human parvovirus B19 |
| Candiru virus segment L | Human adenovirus A | Human papillomavirus 160 | Human picobirnavirus RNA segment 1 |
| Candiru virus segment M | Human adenovirus B1 | Human papillomavirus 161 | Human picobirnavirus RNA segment 2 |
| Candiru virus segment S | Human adenovirus B2 | Human papillomavirus 162 | Human respiratory syncytial virus |
| Canine distemper virus | Human adenovirus C | Human papillomavirus 163 | Human rhinovirus 14 |
| Cell fusing agent virus | Human adenovirus D | Human papillomavirus 164 | Human rhinovirus 89 |
| Chandipura virus isolate CIN 0451 | Human adenovirus E | Human papillomavirus 165 | Human rotavirus B strain Bang373 RNA dependent RNA polymerase (VP1) mRNA complete cds |
| Chaoyang virus | Human adenovirus F | Human papillomavirus 166 | Human rotavirus B strain Bang373 VP3 (VP3) mRNA complete cds |
| Chapare virus segment L | Human astrovirus | Human papillomavirus 169 | Human rotavirus B strain Bang373 inner capsid protein (VP2) gene complete cds |
| Chapare virus segment S | Human bocavirus | Human papillomavirus 17 | Human rotavirus B strain Bang373 inner capsid protein (VP6) gene complete cds |
| Chikungunya virus | Human bocavirus 2 | Human papillomavirus 170 | Human rotavirus B strain Bang373 nonstructural protein (NSP2) gene complete cds |
| Colobus guereza papillomavirus type 2 | Human bocavirus 3 | Human papillomavirus 18 | Human rotavirus B strain Bang373 nonstructural protein (NSP3) gene complete cds |
| Colorado tick fever virus segment 1 | Human bocavirus 4 | Human papillomavirus 19 | Human rotavirus B strain Bang373 nonstructural protein (NSP4) gene complete cds |
| Colorado tick fever virus segment 10 | Human coronavirus 229E | Human papillomavirus 2 | Human rotavirus B strain Bang373 nonstructural protein (NSP5) gene complete cds |
| Colorado tick fever virus segment 11 | Human coronavirus HKU1 | Human papillomavirus 20 | Human rotavirus B strain Bang373 nonstructural protein 1-1 (NSP1-1) nonstructural protein 1-2 (NSP1-2) and nonstructural protein 1-3 (NSP1-3) genes complete cds |
| Colorado tick fever virus segment 12 | Human coronavirus NL63 | Human papillomavirus 21 | Human rotavirus B strain Bang373 outer capsid protein (VP4) gene complete cds |
| Colorado tick fever virus segment 2 | Human coronavirus OC43 | Human papillomavirus 22 | Human rotavirus B strain Bang373 outer capsid protein (VP7) gene complete cds |
| Colorado tick fever virus segment 3 | Human endogenous retrovirus K113 | Human papillomavirus 23 | Ikoma lyssavirus |
| Colorado tick fever virus segment 4 | Human enteric coronavirus strain 4408 | Human papillomavirus 24 | Ilheus virus |
| Colorado tick fever virus segment 5 | Human enterovirus A | Human papillomavirus 25 | Influenza A virus (A/Goose/Guangdong/1/96(H5N1)) |
| Colorado tick fever virus segment 6 | Human enterovirus B | Human papillomavirus 26 | Influenza A virus (A/Goose/Guangdong/1/96(H5N1)) |

### Supplementary Table 1. List of  enrolled viruses (continued)

| | | | |
|---|---|---|---|
| Influenza A virus (A/Goose/Guangdong/1/96(H5N1)) | Louping ill virus | Rotavirus C segment 6 | Torque teno virus 1 |
| Influenza A virus (A/Goose/Guangdong/1/96(H5N1)) | Lujo virus segment L | Rotavirus C segment 7 | Torque teno virus 10 |
| Influenza A virus (A/Goose/Guangdong/1/96(H5N1)) segment 2 | Lujo virus segment S | Rotavirus C segment 8 | Torque teno virus 12 |
| Influenza A virus (A/Goose/Guangdong/1/96(H5N1)) segment 4 | Luna virus segment L | Rotavirus C segment 9 | Torque teno virus 14 |
| Influenza A virus (A/Goose/Guangdong/1/96(H5N1)) strain A/Goose/Guangdong/1/96(H5N1) | Luna virus segment S | Rotavirus F chicken/03V0568/DEU/2003 segment 1 | Torque teno virus 15 |
| Influenza A virus (A/Goose/Guangdong/1/96(H5N1)) strain A/Goose/Guangdong/1/96(H5N1) | Lunk virus NKS-1 segment L | Rotavirus F chicken/03V0568/DEU/2003 segment 10 | Torque teno virus 16 |
| Influenza A virus (A/Hong Kong/1073/99(H9N2)) | Lunk virus NKS-1 segment S | Rotavirus F chicken/03V0568/DEU/2003 segment 11 | Torque teno virus 19 |
| Influenza A virus (A/Hong Kong/1073/99(H9N2)) segment 1 | Lymphocytic choriomeningitis virus segment L | Rotavirus F chicken/03V0568/DEU/2003 segment 2 | Torque teno virus 2 |
| Influenza A virus (A/Hong Kong/1073/99(H9N2)) segment 2 | Lymphocytic choriomeningitis virus segment S | Rotavirus F chicken/03V0568/DEU/2003 segment 3 | Torque teno virus 25 |
| Influenza A virus (A/Hong Kong/1073/99(H9N2)) segment 3 | Macaca fascicularis papillomavirus type 2 | Rotavirus F chicken/03V0568/DEU/2003 segment 4 | Torque teno virus 26 |
| Influenza A virus (A/Hong Kong/1073/99(H9N2)) segment 4 | Machupo virus segment L | Rotavirus F chicken/03V0568/DEU/2003 segment 5 | Torque teno virus 27 |
| Influenza A virus (A/Hong Kong/1073/99(H9N2)) segment 6 | Machupo virus segment S | Rotavirus F chicken/03V0568/DEU/2003 segment 6 | Torque teno virus 28 |
| Influenza A virus (A/Hong Kong/1073/99(H9N2)) segment 7 | Mapuera virus | Rotavirus F chicken/03V0568/DEU/2003 segment 7 | Torque teno virus 3 |
| Influenza A virus (A/Hong Kong/1073/99(H9N2)) segment 8 | Mayaro virus | Rotavirus F chicken/03V0568/DEU/2003 segment 8 | Torque teno virus 4 |
| Influenza A virus (A/Korea/426/1968(H2N2)) | Measles virus | Rotavirus F chicken/03V0568/DEU/2003 segment 9 | Torque teno virus 6 |
| Influenza A virus (A/Korea/426/68(H2N2)) segment 2 | Menangle virus | Rotavirus G chicken/03V0567/DEU/2003 segment 1 | Torque teno virus 7 |
| Influenza A virus (A/Korea/426/68(H2N2)) segment 3 | Merkel cell polyomavirus | Rotavirus G chicken/03V0567/DEU/2003 segment 10 | Torque teno virus 8 |
| Influenza A virus (A/Korea/426/68(H2N2)) segment 4 | Mobala virus segment L | Rotavirus G chicken/03V0567/DEU/2003 segment 11 | Toscana virus segment L |
| Influenza A virus (A/Korea/426/68(H2N2)) segment 5 | Mobala virus segment S | Rotavirus G chicken/03V0567/DEU/2003 segment 2 | Toscana virus segment M |
| Influenza A virus (A/Korea/426/68(H2N2)) segment 6 | Modoc virus | Rotavirus G chicken/03V0567/DEU/2003 segment 3 | Toscana virus segment S |
| Influenza A virus (A/Korea/426/68(H2N2)) segment 7 | Mokola virus | Rotavirus G chicken/03V0567/DEU/2003 segment 4 | Tula virus segment L |
| Influenza A virus (A/Korea/426/68(H2N2)) segment 8 | Moloney murine leukemia virus | Rotavirus G chicken/03V0567/DEU/2003 segment 5 | Tula virus segment M |
| Influenza A virus (A/New York/392/2004(H3N2)) segment 1 | Montana myotis leukoencephalitis virus | Rotavirus G chicken/03V0567/DEU/2003 segment 6 | Tula virus segment S |
| Influenza A virus (A/New York/392/2004(H3N2)) segment 2 | Mopeia Lassa reassortant 29 segment L | Rotavirus G chicken/03V0567/DEU/2003 segment 7 | Tupaia paramyxovirus |
| Influenza A virus (A/New York/392/2004(H3N2)) segment 4 | Mopeia Lassa reassortant 29 segment S | Rotavirus G chicken/03V0567/DEU/2003 segment 8 | Usutu virus |
| Influenza A virus (A/New York/392/2004(H3N2)) segment 5 | Mopeia virus AN20410 segment L | Rotavirus G chicken/03V0567/DEU/2003 segment 9 | Uukuniemi virus |
| Influenza A virus (A/New York/392/2004(H3N2)) segment 6 | Mopeia virus AN20410 segment S | Rubella virus | Uukuniemi virus chromosome segment M |
| Influenza A virus (A/New York/392/2004(H3N2)) segment 7 | Morogoro virus segment L | SARS coronavirus | Uukuniemi virus segment L |
| Influenza A virus (A/New York/392/2004(H3N2)) segment 8 | Morogoro virus segment S | SFTS virus HB29 segment L | Vaccinia virus |
| Influenza A virus (A/New York/392/2004(H3N2)) strain A/New York/392/2004 | Mosquito flavivirus isolate LSFlaviV-A20-09 | SFTS virus HB29 segment M | Variola virus |
| Influenza A virus (A/Puerto Rico/8/34(H1N1)) segment 1 | Mossman virus | SFTS virus HB29 segment S | Vesicular stomatitis Indiana virus |
| Influenza A virus (A/Puerto Rico/8/34(H1N1)) segment 2 | Mouse mammary tumor virus | Sabia virus | WU Polyomavirus |
| Influenza A virus (A/Puerto Rico/8/34(H1N1)) segment 3 | MuLV_DG75 | Sabia virus segment L | Wesselsbron virus |
| Influenza A virus (A/Puerto Rico/8/34(H1N1)) segment 4 | MuLV_JY | Sandfly Sicilian Turkey virus segment L | West Nile virus |
| Influenza A virus (A/Puerto Rico/8/34(H1N1)) segment 5 | MuLV_MCF1233 | Sandfly Sicilian Turkey virus segment M | West Nile virus |
| Influenza A virus (A/Puerto Rico/8/34(H1N1)) segment 6 | MuLV_N417 | Sandfly Sicilian Turkey virus segment S | Whitewater Arroyo virus segment L |
| Influenza A virus (A/Puerto Rico/8/34(H1N1)) segment 7 | Mumps virus | Sapovirus C12 strain C12 | Whitewater Arroyo virus segment S |
| Influenza A virus (A/Puerto Rico/8/34(H1N1)) segment 8 | Murine type C retrovirus | Sapovirus Hu/Dresden/pJG-Sap01/DE | XMRV_VP62 |
| Influenza B virus RNA 4 | Murray Valley encephalitis virus | Sapovirus Mc10 | Yellow fever virus |
| Influenza B virus RNA 5 | Nariva virus | Sathuperi virus Gn-Gc-NSm gene for M polyprotein segment M genomic RNA | Yokose virus |
| Influenza B virus RNA 6 | Newcastle disease virus B1 | Sathuperi virus N and NSs genes segment S genomic RNA | Zika virus |
| Influenza B virus RNA 7 | Nipah virus | Sathuperi virus RdRp gene for RNA-dependent RNA polymerase segment L genomic RNA |  |
| Influenza B virus RNA 8 | Norwalk virus | Seal anellovirus TFFN/USA/2006 | |
| Influenza B virus RNA-2 | Ntaya virus isolate IPDIA | Sendai virus | |
| Influenza B virus RNA-3 | O nyong-nyong virus | Seoul virus segment M | |
| Influenza C virus (C/Ann Arbor/1/50) segment 1 partial sequence | Oliveros virus segment L | Seoul virus strain 80-39 segment S |  |
| Influenza C virus (C/Ann Arbor/1/50) segment 2 | Oliveros virus segment S | Seoul virus strain Seoul 80-39 clone 1 | |
| Influenza C virus (C/Ann Arbor/1/50) segment 3 | Omsk hemorrhagic fever virus | Sepik virus | |
| Influenza C virus (C/Ann Arbor/1/50) segment 4 | Oropouche virus segment L | Severe acute respiratory syndrome coronavirus 2 isolate Wuhan-Hu-1 | |
| Influenza C virus (C/Ann Arbor/1/50) segment 5 | Oropouche virus segment M | Shamonda virus Gn-Gc-NSm gene for M polyprotein segment M genomic RNA isolate Ib An 5550 |  |
| Influenza C virus (C/Ann Arbor/1/50) segment 6 | Oropouche virus segment S | Shamonda virus N and NSs genes segment S genomic RNA isolate Ib An 5550 |  |
| Influenza C virus (C/Ann Arbor/1/50) segment 7 | Parainfluenza virus 5 | Shamonda virus RdRp gene for RNA-dependent RNA polymerase segment L genomic RNA isolate Ib An 5550 |  |
| Ippy virus segment L | Parana virus segment L | Simbu virus Gn-Gc-NSm gene for M polyprotein segment M genomic RNA isolate SA Ar 53 |  |
| Ippy virus segment S | Parana virus segment S (small) | Simbu virus N and NSs genes segment S genomic RNA isolate SA Ar 53 |  |
| Irkut virus | Peste-des-petits-ruminants virus | Simbu virus RdRp gene for RNA-dependent RNA polymerase segment L genomic RNA isolate SA Ar 53 |  |
| J-virus | Pichinde virus | Simian virus 40 | |
| JC polyomavirus | Pichinde virus L RNA | Simian virus 41 | |
| Japanese encephalitis virus genome | Pirital virus segment L | Sin Nombre virus segment L |  |
| Junin virus segment L | Pirital virus segment S | Sin Nombre virus segment M | |
| Junin virus segment S | Pneumonia virus of mice J3666 | Sin Nombre virus segment S | |
| KI polyomavirus Stockholm 60 | Poliovirus | Small anellovirus 1 | |
| Kadipiro virus chromosome segment 1 | Porcine enteric sapovirus | Small anellovirus 2 | |
| Kadipiro virus chromosome segment 10 | Porcine rubulavirus | St. Louis encephalitis virus | |
| Kadipiro virus chromosome segment 12 | Powassan virus | Sudan ebolavirus | |
| Kadipiro virus chromosome segment 2 | Puumala virus segment L | TTV-like mini virus isolate TTMV LY1 | |
| Kadipiro virus chromosome segment 3 | Puumala virus segment M | Tacaribe virus segment L | |
| Kadipiro virus chromosome segment 4 | Puumala virus segment S | Tacaribe virus segment S | |
| Kadipiro virus chromosome segment 5 | Quang Binh virus | Tamana bat virus genome |  |
| Kadipiro virus chromosome segment 6 | Rabies virus | Tamiami virus segment L | |
| Kadipiro virus chromosome segment 8 | Rauscher murine leukemia virus | Tamiami virus segment S (small) | |
| Kadipiro virus chromosome segment 9 | Razdan virus strain LEIV-Arm2741 segment L | Tembusu virus strain JS804 | |
| Kadipiro virus segment 11 | Razdan virus strain LEIV-Arm2741 segment M | Thogoto virus | |
| Kadipiro virus segment 7 | Razdan virus strain LEIV-Arm2741 segment S | Thogoto virus | |
| Kamiti River virus | Respiratory syncytial virus | Thogoto virus | |
| Karshi virus | Reston ebolavirus | Thogoto virus segment 1 | |
| Kedougou virus | Rift Valley fever virus segment L | Thogoto virus segment 5 | |
| Kokobera virus | Rift Valley fever virus segment M | Thogoto virus segment 6 | |
| La Crosse virus segment L | Rift Valley fever virus segment S | Thottapalayam virus segment L | |
| La Crosse virus segment M | Rinderpest virus (strain Kabete O) | Thottapalayam virus segment M | |
| La Crosse virus segment S | Rio Bravo virus genome | Thottapalayam virus segment S |  |
| Lagos bat virus isolate 0406SEN | Rodent hepacivirus isolate RHV-339 | Tick-borne encephalitis virus | |
| Lake Victoria marburgvirus - Musoke | Ross River virus | Tioman virus | |
| Langat virus | Rotavirus A segment 1 | Torque teno canis virus | |
| Lassa virus segment L | Rotavirus A segment 10 | Torque teno douroucouli virus | |
| Lassa virus segment S | Rotavirus A segment 11 | Torque teno felis virus | |
| Latino virus segment L | Rotavirus A segment 2 | Torque teno midi virus 1 | |
| Latino virus segment S | Rotavirus A segment 3 | Torque teno midi virus 2 | |
| Liao ning virus segment 1 | Rotavirus A segment 4 | Torque teno mini virus 1 | |
| Liao ning virus segment 10 | Rotavirus A segment 5 | Torque teno mini virus 2 | |
| Liao ning virus segment 11 | Rotavirus A segment 6 | Torque teno mini virus 3 | |
| Liao ning virus segment 12 | Rotavirus A segment 7 | Torque teno mini virus 4 | |
| Liao ning virus segment 2 | Rotavirus A segment 8 | Torque teno mini virus 5 | |
| Liao ning virus segment 3 | Rotavirus A segment 9 | Torque teno mini virus 6 | |
| Liao ning virus segment 4 | Rotavirus C segment 1 | Torque teno mini virus 7 | |
| Liao ning virus segment 5 | Rotavirus C segment 10 | Torque teno mini virus 8 | |
| Liao ning virus segment 6 | Rotavirus C segment 11 | Torque teno mini virus 9 | |
| Liao ning virus segment 7 | Rotavirus C segment 2 | Torque teno sus virus 1 | |
| Liao ning virus segment 8 | Rotavirus C segment 3 | Torque teno sus virus k2 isolate 2p | |
| Liao ning virus segment 9 | Rotavirus C segment 4 | Torque teno tamarin virus | |
| Lloviu virus | Rotavirus C segment 5 | Torque teno virus | |

Supplementary Table 1. The full list of enrolled viruses in VIRTUS.
